# What makes a society dead: accumulation of uric acid increases infectious disease risk in termites

**DOI:** 10.1101/2025.04.23.650172

**Authors:** Takao Konishi, Masaaki Nakashima, Tomonari Nozaki, Eisuke Tasaki, Mamoru Takata, Kenji Matsuura

**Affiliations:** Laboratory of Insect Ecology, Graduate School of Agriculture, Kyoto University, Kitashirakawa-Oiwake-cho, Sakyo-ku, Kyoto 606-8502, Japan; Department of Forest Entomology, Forestry and Forest Products Research Institute, 1 Matsunosato, Tsukuba, Ibaraki 305-8687, Japan; Laboratory of Evolutionary Genomics, National Institute for Basic Biology, Okazaki, Aichi 444-8585, Japan; Department of Biology, Faculty of Science, Niigata University, 8050 Ikarashi 2-no-cho, Nishi-ku, Niigata 950-2181, Japan

**Keywords:** colony collapse, eusocial insect, opportunistic infection, social immunity, urate

## Abstract

In social insect colonies, the high reproductive output of kings and queens is supported by non-reproductive castes, a division of labour that allows for the rapid recovery of individuals, often including those of the reproductive castes. However, these seemingly permanent colonies eventually face collapse. Studying the mechanisms of colony death is crucial for understanding the maintenance of insect social systems. Here, we show that the accumulation of uric acid (a major product of nitrogen metabolism) in workers increases their infectious disease risk in the Japanese subterranean termite *Reticulitermes speratus*. In this species, king replacement has been associated with colony demise, and we found that king replacement increases the uric acid contents in worker bodies in the field. Uric acid, which has antioxidant activity, was shown to reduce *in vivo* levels of reactive oxygen species (ROS), a key player in innate immunity. Workers with decreased ROS levels were more susceptible than those with normal ROS levels to an opportunistic pathogen that causes disease to immunocompromised termites. Our results indicate that regulating the individual oxidant/antioxidant balance through the interactions among colony members plays a pivotal role in the immunity of social insects.

## 1. Introduction

Social insects, such as termites, ants, bees and wasps, are the most abundant animals on earth [1–3]. Their ecological success is attributed to the division of labour within their colonies that consist of many, typically related individuals. They have separated the roles of reproductive castes (kings and queens) and non-reproductive castes (workers and soldiers), and increased their fecundity by specialising in each task [4,5]. Reproductive castes of social insects continuously exhibit enormous reproductive output, supported by other castes engaging in non-reproductive tasks such as foraging and defence [6,7]. For example, in the termite species *Macrotermes subhyalinus*, a queen is estimated to lay approximately 40,000 eggs (corresponding to about one-third of its fresh weight) per day [8]. Such high fecundity allows for the rapid recovery of accidentally lost individuals, often including those of the reproductive caste [9,10]. Although such colonies appear permanent, they eventually collapse. Studying the mechanisms involved in colony death is crucial for understanding how social systems in insects are maintained; at present, the molecular and physiological bases are largely unknown.

Colonies of social insects face various extrinsic mortality factors such as predation, disease, starvation, desiccation and conflict with neighbouring colonies [11–13]. However, because the long-term fate of each colony is unique depending on the environmental threats [14,15], colony death is a poorly studied aspect of the life of social insects, except for colony collapse disorder (CCD) in honeybees. The decline of honeybee colonies and their eventual collapse has been widespread throughout the Northern Hemisphere and this phenomenon is a matter of concern for both beekeepers and scientists [16]. Recent studies have provided clues on the loss of colony functions; this dramatic event is associated with an enhanced impact of parasites and pathogens on honeybees, which is indicative of reduced immunocompetence [17,18]. The increase of opportunistic microbes and arthropods in dying colonies was also reported in the termite *Coptotermes formosanus* [19]. However, beyond such observations, information on the mechanisms of colony death in social insects remains fragmentary. Another social insect model is needed for a more comprehensive and integrated analysis.

The Japanese subterranean termite *Reticulitermes speratus* is one of the most well-studied social insect species in terms of molecular and physiological analysis due to its unique reproductive system, named asexual queen succession (AQS) and extraordinary longevity of reproductive castes [7,20]. Similar to other termite species, *R. speratus* colonies are founded by a monogamous pair of primary reproductives derived from alates (winged adults). In AQS, the primary queen is genetically immortal until a colony dies due to the production of a large number of secondary queens through parthenogenesis [21]. This system enables queen succession in the colony without inbreeding. In contrast to the queen, a primary king should continue to live even if queens are replaced, because the loss of a primary king and subsequent emergence of secondary kings (sons of the primary king and primary/secondary queens) will result in mother-son inbreeding (figure 1). It is true that colonies headed by secondary kings are rare in the field [22], while how king replacement causes the colony’s end has never been empirically demonstrated.

**Figure 1.**
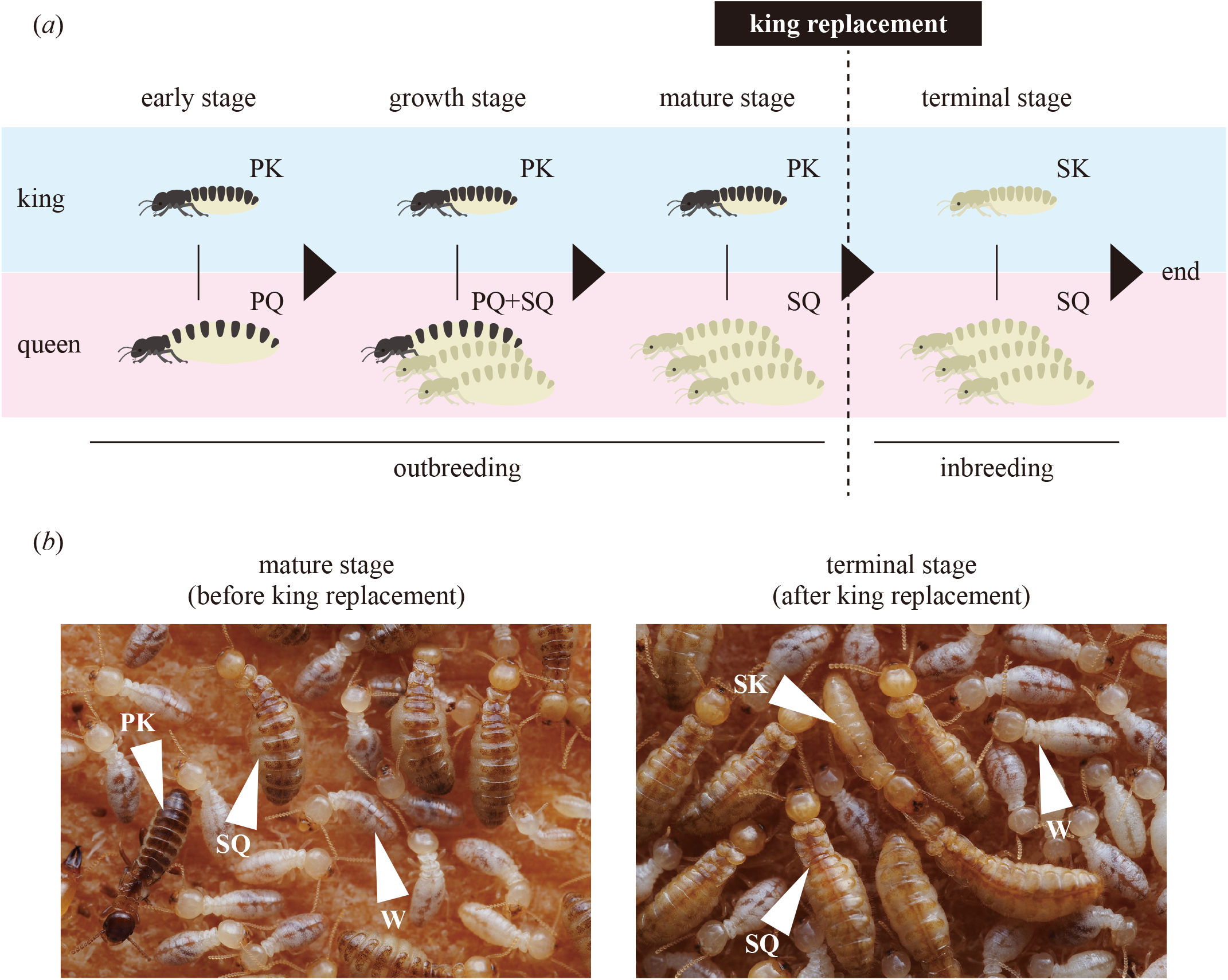
Colony collapse associated with king replacement in the subterranean termite *Reticulitermes speratus*. (*a*) A simplified schematic model of king and queen succession in *R. speratus*. In this species, the loss of a primary king and subsequent emergence of a secondary king has been associated with colony demise because colonies after king replacement are rare in the field. (*b*) Representative photograph of termite colonies before (left panels) and after (right panels) king replacement. PK, primary king; PQ, primary queen; SK, secondary king; SQ, secondary queen; W, worker. (Online version in colour.)

Konishi et al. [23] found a potential clue to tackle this conundrum: under laboratory conditions, the removal of primary kings from *R. speratus* colonies leads to the accumulation of uric acid (a major product of nitrogen metabolism) in worker bodies. Uric acid is a strong antioxidant against reactive oxygen species (ROS) in organisms [24]. ROS are oxygen-derived radical species (e.g. superoxide, hydroxyl radical, hydrogen peroxide and singlet oxygen) that are continuously formed as byproducts of aerobic metabolism. Excessive ROS generation can damage biomolecules, such as DNA, proteins and lipids, through oxidative stress; however, appropriate amounts of ROS is beneficial to immune defence in animals and plants [25–27]. Therefore, the accumulation of uric acid can deplete ROS in termites and compromise their immune system. A comparative analysis focusing on uric acid between colonies before and after king replacement in *R. speratus* can provide an excellent opportunity for investigating mechanisms of colony death in social insects.

In this study, we first conducted a field survey to compare the uric acid levels in workers before and after king replacement, which has been associated with colony demise in *R. speratus*. Because we found that uric acid accumulates accompanied by king replacement also in the field, we examined the negative effect of uric acid accumulation in termites. We confirmed that the accumulation of uric acid, which has antioxidant activity, reduces *in vivo* ROS, a key player in innate immunity. We then investigated the survival rates of workers with increased levels of uric acid when exposed to an opportunistic pathogen, which causes disease in immunocompromised hosts. Finally, to demonstrate the influences of ROS contents on infectious disease risk, we performed administration of another antioxidant.

## 2. Materials and Methods

### (a) Samples and treatments

We collected colonies of *R. speratus* along with nest wood from secondary forests in Japan. As with other subterranean termites, *R. speratus* has cryptic nesting habits and builds transient, hidden royal chambers deep inside wood. To find colonies, we first searched for termite workers in decayed logs on the ground. Then, we located the royal chamber area based on the distribution of eggs and 1st instar larvae. We obtained the parts of the wood containing the royal chambers using a saw and brought them into the laboratory for further dismantling. To avoid counting supplemental reproductives that differentiated after collection, all kings and queens were extracted from the logs within 10 days of collection. We distinguished primary kings/queens (alate-derived) from secondary kings/queens (neotenic) based on the formers’ fully melanised body colour and the presence of wing bases. We distinguished the sex of each caste by the morphology of the caudal sternites, as described in a previous study [28]. All colonies were maintained under constant darkness at 25°C in the laboratory and fed brown-rotted pinewood mixed cellulose (BPC) medium [29]. No specific permits or licences were required for this study. We followed the Kyoto University Regulations on Animal Experimentation for the treatment of animals.

First, we measured the uric acid levels in workers before and after the king replacement, which has been associated with colony demise in *R. speratus*. In termites, uric acid is stored in specialised cells of fat bodies that perform liver-like functions [30–32]. As reported in previous studies [30–32], the fat bodies of workers consisted of two types of cells: ‘adipocytes’ containing lipid droplets and ‘urocytes’ containing urate crystals (figure 2*a*). Because we observed the increase of urate crystals in the fat bodies of workers accompanied by king replacement, we additionally compared the uric acid contents before and after king replacement to quantitatively evaluate the uric acid accumulation. We randomly chose five male and five female workers and measured the uric acid contents. We processed a total of 10 colony replicates for each treatment.

**Figure 2.**
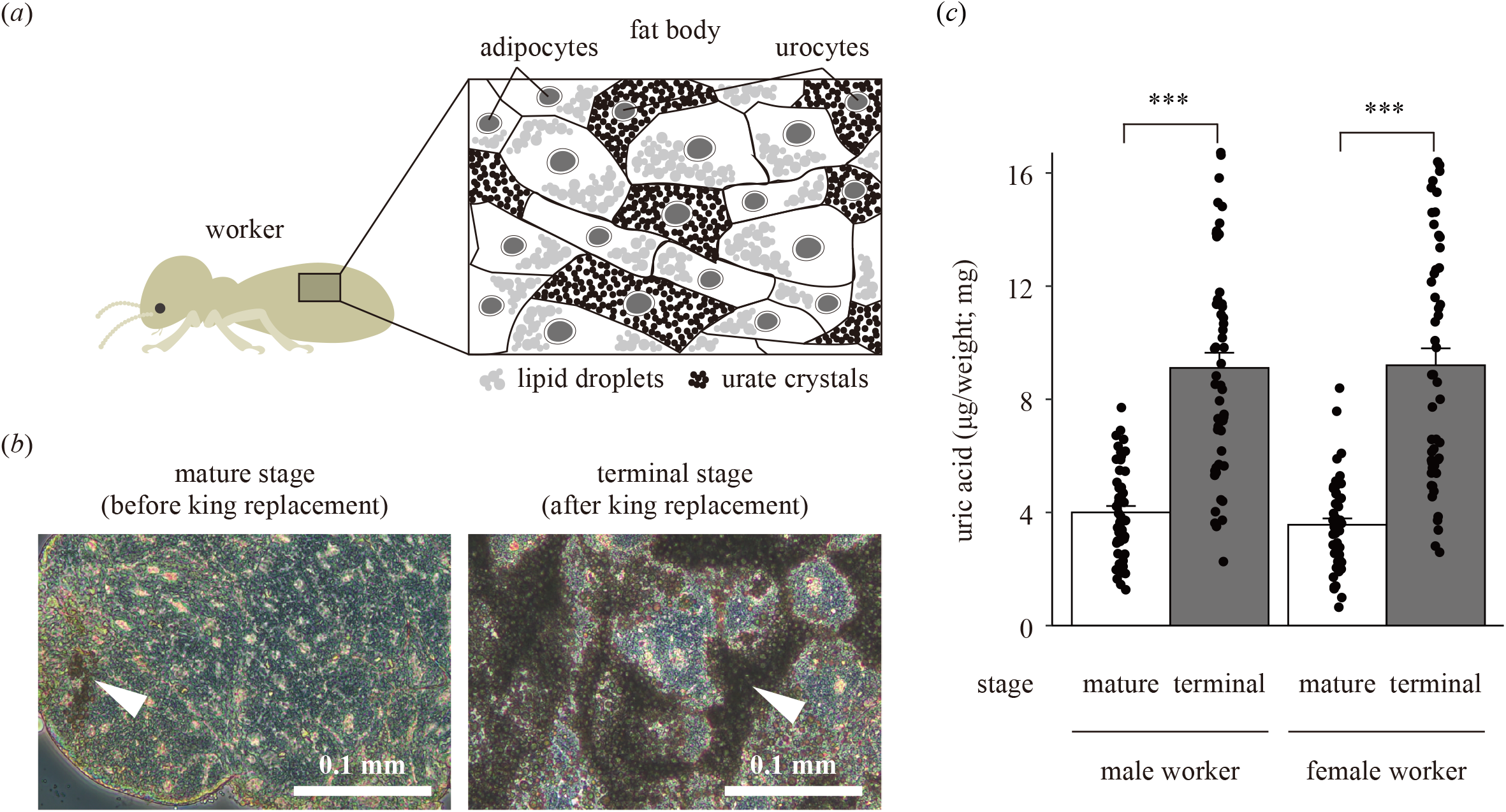
Comparison of uric acid accumulation in *Reticulitermes speratus* workers in colonies before and after king replacement. (*a*) Fat body tissue of termite workers composed of two types of cells: adipocytes containing lipid droplets and urocytes with urate crystals. (*b*) Phase-contrast microscopy comparing the morphologies of worker fat bodies in colonies before (left panels) and after (right panels) king replacement. White arrows indicate urate crystals accumulated in urocytes. (*c*) Comparison of the uric acid contents of workers before and after king replacement. Black points show individual termites, jittered to reduce overlap between points. Error bars denote the standard error of the mean (SEM). Asterisks indicate significant differences (likelihood ratio test, \**p* < 0.05, \*\**p* < 0.01, \*\*\**p* < 0.001). (Online version in colour.)

Next, to examine the effects of uric acid (which has antioxidant activity) on the levels of ROS in worker bodies, we established the following two groups: UA+ and UA−. Specifically, the UA+ group consisted of 100 workers that were placed in a 40-mm Petri dish filled with 3 g BPC medium [29], with uric acid (FUJIFILM Wako Pure Chemical, Osaka, Japan) at a final concentration of 1% (w/w). The UA− group consisted of 100 workers similarly placed in a Petri dish filled with BPC medium only. We randomly extracted four male and four female workers and compared the uric acid contents (two males and two females) and ROS contents (two males and two females) between the two groups prepared by the above methods after 1, 2 and 3 weeks. We used termites from four colonies to conduct the experiments. We visualised the changes in the fresh fat bodies of workers due to the accumulation of uric acid by phase-contrast microscopy.

Finally, to investigate the effects of uric acid accumulation on the immune function in termites, we compared the survival rates of workers when exposed to the opportunistic pathogen, which causes disease in immunocompromised hosts. We used 3-week workers of the two groups for the infection assays because the increase of uric acid contents and decrease of ROS contents were observed in the feeding experiments. As an opportunistic pathogen, we used *Serratia marcescens* isolated from the nest of *R. speratus. S. marcescens* is a typical opportunistic bacterium that is ubiquitous in the environment and is regularly isolated from the nests and corpses of termites [34–36]. This bacterium generally does not harm healthy termites but causes disease to immunocompromised hosts [37]. A red pigment called prodigiosin produced by *S. marcescens* enables us to easily detect this species [38]. *S. marcescens* lacks urate oxidase, which is required to metabolise uric acid [39,40]. In the infection assays, 30 workers were held in a Petri dish (40 mm in diameter) lined with filter paper (35 mm in diameter) containing 200 µL of the bacterial suspension (4.3 × 10^10^ CFUs/mL). As a control, 30 workers were held in a Petri dish (40 mm in diameter) lined with filter paper (35 mm in diameter) containing 200 µL of the lysogeny broth (LB) liquid media only. The colony size (30 individuals) exceeds the minimum to maintain for longer than one year under laboratory conditions reported in previous studies using *Reticulitermes* termites [41]. After the *Serratia* exposure treatment, we recorded the survival rate for ten days. In addition, to demonstrate the influences of ROS on the infectious disease risk, we performed feeding of another antioxidant L-ascorbic acid (FUJIFILM Wako Pure Chemical, Osaka, Japan) under the same conditions as the above method of uric acid administration.

### (b) Microscopic observation of fat bodies

To observe the changes in fat bodies associated with the accumulation of uric acid, we performed a morphological analysis of fresh fat bodies of *R. speratus* workers using a microscope. The fat bodies were dissected in phosphate-buffered saline (PBS; 33 mM KH_2_PO_4_ and 33 mM Na_2_HPO_4_, pH 6.8) using fine forceps under a stereomicroscope (Olympus SZX7; Olympus, Tokyo, Japan). We dissected the fat bodies, which appear as whitish loose tissues located around digestive tubes and reproductive organs, from the abdomens of the insects with care to avoid contamination by other tissues, such as Malpighian tubules and tracheoles. The dissected tissues were directly mounted on glass slides and fixed with 4% paraformaldehyde (PFA) in PBS for 5 min, crushed using coverslips and observed using a phase-contrast microscope (Leica DM IL LED; Leica, Hamburg, Germany).

### (c) Quantification of uric acid content

We measured uric acid contents using a Uric Acid Assay Kit (Abcam, Cambridge, UK) according to the manufacturer’s protocols. Briefly, each whole termite body was homogenised in 100 μL of Uric Acid Assay Buffer and centrifuged at 15,000 rpm for 2 min at 4°C. Supernatants were collected as sample extracts and diluted 10-fold. 50 μL of the diluted sample extract was mixed with 50 μL of the Fluorometric Reaction Mix (0.4 μL Uric Acid Probe, 2.0 μL Uric Acid Enzyme Mix, and 47.6 μL Uric Acid Assay Buffer) and incubated for 30 min at 37°C in the dark. The uric acid content was determined fluorometrically (Ex/Em = 530/590 nm) using a Fluoroskan Ascent FL microplate reader (Thermo Fisher Scientific, Waltham, MA, USA) alonh with serially diluted uric acid standards (0, 0.8, 1.6, 2.4, 3.2 and 4.0 nmol/well). To eliminate differences in body size, the uric acid content was standardised by dividing by the fresh body weight.

### (d) Quantification of ROS content

ROS contents were measured using a ROS Assay Kit (Dojindo, Kumamoto, Japan) according to the manufacturer’s protocols. Briefly, each sample was homogenised in 150 μL of Highly Sensitive DCFH-DA Working Solution (0.15 μL Highly Sensitive DCFH-DA Dye and 149.85 μL Loading Buffer) and kept for 30 min at room temperature (25°C). After centrifugation at 15,000 rpm for 2 min at 25°C, 100 μL of the supernatant was collected. The ROS content was determined fluorometrically (Ex/Em = 485/538 nm) using a Fluoroskan Ascent FL microplate reader (Thermo Fisher Scientific). The measured fluorescence intensity was expressed as the relative ROS content in arbitrary units (AU). To eliminate differences in body size, the ROS content was standardised by dividing by the fresh body weight.

### (e) Isolation and culture of *S. marcescens*

For the infection assays, *S. marcescens* was isolated from the nest of *R. speratus*. We put the dead body turned red on the LB agar plate and incubated it at 28°C for 24 h. A red-pigmented colony was picked up and streaked out on fresh LB agar medium using sterile platinum loops to purify the isolate. A bacterial suspension was prepared by transferring a colony of *S. marcescens* into LB liquid medium and incubating at 28°C for 24 h. We identified the species of the bacterium by sequencing the 16S rRNA gene. The genomic DNA used for amplification was extracted from the bacteria in LB liquid culture and purified as previously described [41]. The 16S rRNA gene was amplified using the 10F/800R primers (Forward: 5’-GTTTGATCCTGGCTCA-3’, Reverse: 5’-TACCAGGGTATCTAATCC-3’). The purified PCR products were sequenced using an ABI 3500 Genetic Analyzer (Applied Biosystems, Foster City, CA, USA). We compared the resultant sequence (565 bp) with those in GenBank using the blastn programme [42]. This bacterium showed 100% homology with a published 16S rRNA gene sequence from *S. marcescens* strain KABOSH1 (GenBank accession number: OQ550114). To determine the bacterial concentration, serially diluted suspensions were put in LB agar plates, and the colony-forming units (CFUs) were counted after a 24-h incubation.

### (f) Data analysis

All statistical analyses were performed using R software (v4.4.2) [43]. We analysed the uric acid and ROS contents using generalised linear mixed models (GLMMs) with a Gaussian distribution. In the GLMM, treatment was a fixed factor, and colony was a random factor. Likelihood ratio tests were conducted to determine the significance of each fixed effect. A significance value of *p* < 0.05 was considered to indicate statistical significance. In addition, we analysed the survival rates using log-rank tests with Bonferroni corrections. In the log-rank test, treatment was a fixed factor. A significance value of *p* < 0.008 (i.e. 0.05/6) was considered to indicate statistical significance. Experimental data analysed during this study are included in the Supplementary Information file (Dataset S1).

## 3. Results

### (a) Comparative analysis of colonies before and after king replacement

The uric acid contents of worker bodies increased after the king was replaced in the field. In the morphological analysis using a phase-contrast microscope, the increase of urate crystals in the fat bodies was observed in the colonies after king replacement (figure 2*b*). The uric acid contents of workers in colonies after king replacement (headed by secondary kings and secondary queens; terminal stage) were significantly higher than those in colonies before king replacement (headed by primary kings and secondary queens; mature stage) (likelihood ratio test, male: df = 1, *χ*^*2*^ = 17.48, *p <* 0.001; female: df = 1, *χ*^*2*^ = 17.57, *p <* 0.001, figure 2*c*). This trend was seen for both male and female workers.

### (b) Effects of uric acid accumulation on *in vivo* ROS

Feeding of uric acid increased the uric acid contents and decreased the ROS contents in worker bodies. In the morphological analysis using a phase-contrast microscope, the increase of urate crystals in fat bodies was observed during the 3-week feeding of uric acid (figure 3*a*). The uric acid contents of workers in the UA+ groups were significantly higher than those in the UA− groups (likelihood ratio test, week 1: df = 1, *χ* ^*2*^ = 103.59, *p <* 0.001; week 2: df = 1, *χ*^*2*^ = 173.40, *p <* 0.001; week 3: df = 1, *χ*^*2*^ = 303.92, *p <* 0.001, figure 3*b*). In contrast, the ROS contents of workers in the UA+ groups were significantly lower than those in the UA− groups (likelihood ratio test, week 1: df = 1, *χ*^*2*^ = 8.72, *p* = 0.005; week 2: df = 1, *χ*^*2*^ = 6.22, *p* = 0.016; week 3: df = 1, *χ*^*2*^ = 8.80, *p* = 0.005, figure 3*c*). No significant differences in the survival rates were observed between the above two groups during the feeding experiments (likelihood ratio test, week 1: df = 1, *χ*^2^ = 0.02, *p* = 0.895; week 2: df = 1, *χ*^2^ = 0.54, *p* = 0.462; week 3: df = 1, *χ*^2^ = 0.17, *p* = 0.682).

**Figure 3.**
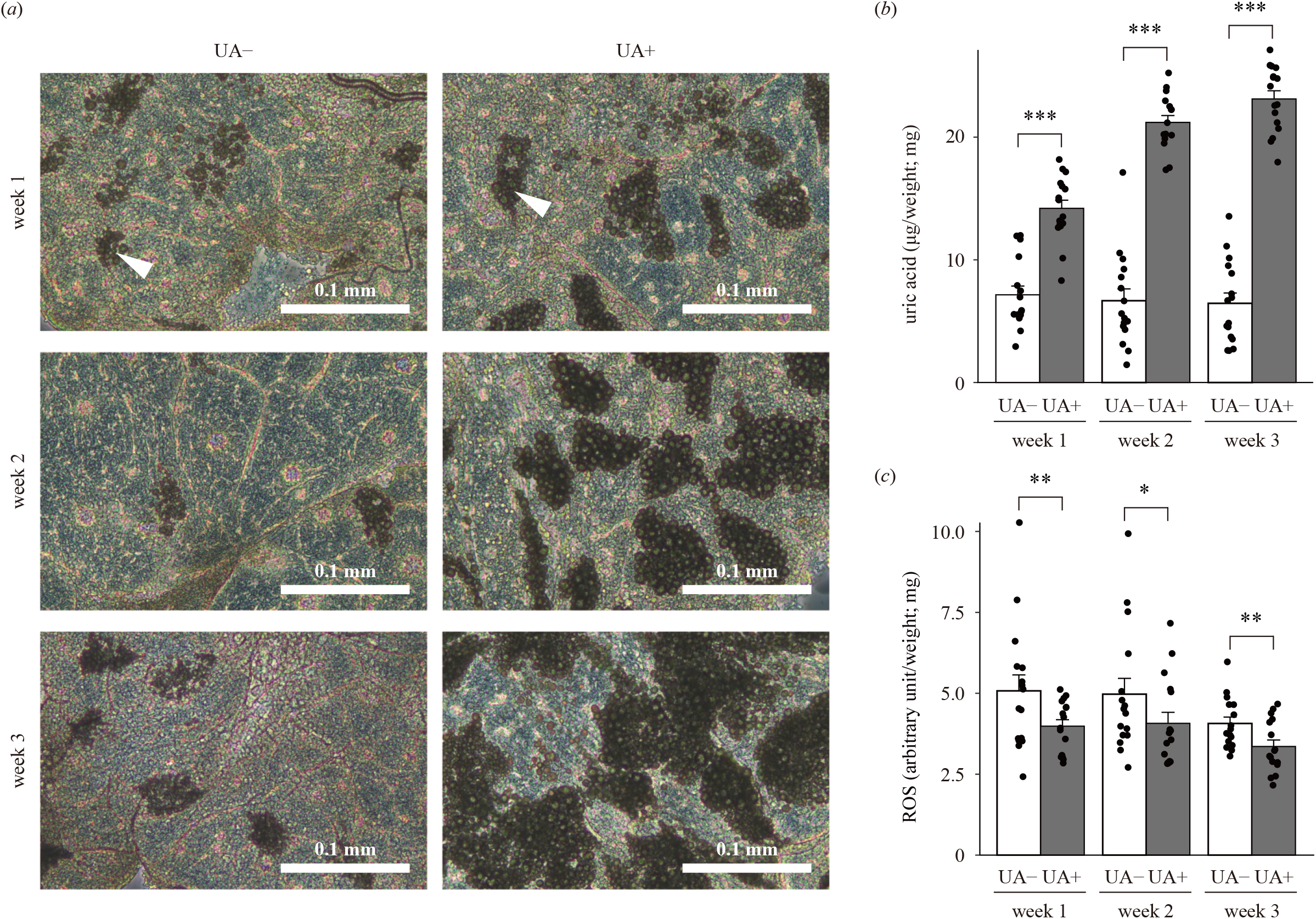
Effects of uric acid accumulation on reactive oxygen species (ROS) in worker bodies. (*a*) Phase-contrast microscopy of uric acid accumulation in fat bodies. White arrows indicate urate crystals accumulated in fat body cells. Comparison of (*b*) uric acid contents and (*c*) ROS contents of workers fed uric acid mixed in brown-rotted pinewood mixed cellulose (BPC) medium and those fed BPC medium only (‘UA+’ and ‘UA−’, respectively). Black points show individual termites, jittered to reduce overlap between points. Error bars denote the standard error of the mean (SEM). Asterisks indicate significant differences (likelihood ratio test, \**p* < 0.05, \*\**p* < 0.01, \*\*\**p* < 0.001). (Online version in colour.)

### (c) Infection assays using opportunistic bacteria

Intriguingly, the accumulation of uric acid increased infectious disease risk in termites. In the infection assays, we observed termite bodies turning red as they died from opportunistic infection by *S. marcescens* (figure 4*a*). Workers with increased uric acid contents exhibited significantly lower viability against the *Serratia* exposure treatment (log-rank test with Bonferroni correction, *p* < 0.001, figure 4*b*). In control (no infection), no significant differences in the survival rates were observed between the two groups during the infection assays (log-rank test with Bonferroni correction, *p* = 1.000, figure 4*b*). Additionally, workers fed another antioxidant (ascorbic acid) also exhibited significantly lower viability against the *Serratia* exposure treatment (log-rank test with Bonferroni correction, *p* < 0.001, figure 4*c*). We also confirmed that feeding ascorbic acid decreased the ROS contents in worker bodies (likelihood ratio test, df = 1, *χ*^2^ = 4.42, *p* = 0.039, figure S1).

**Figure 4.**
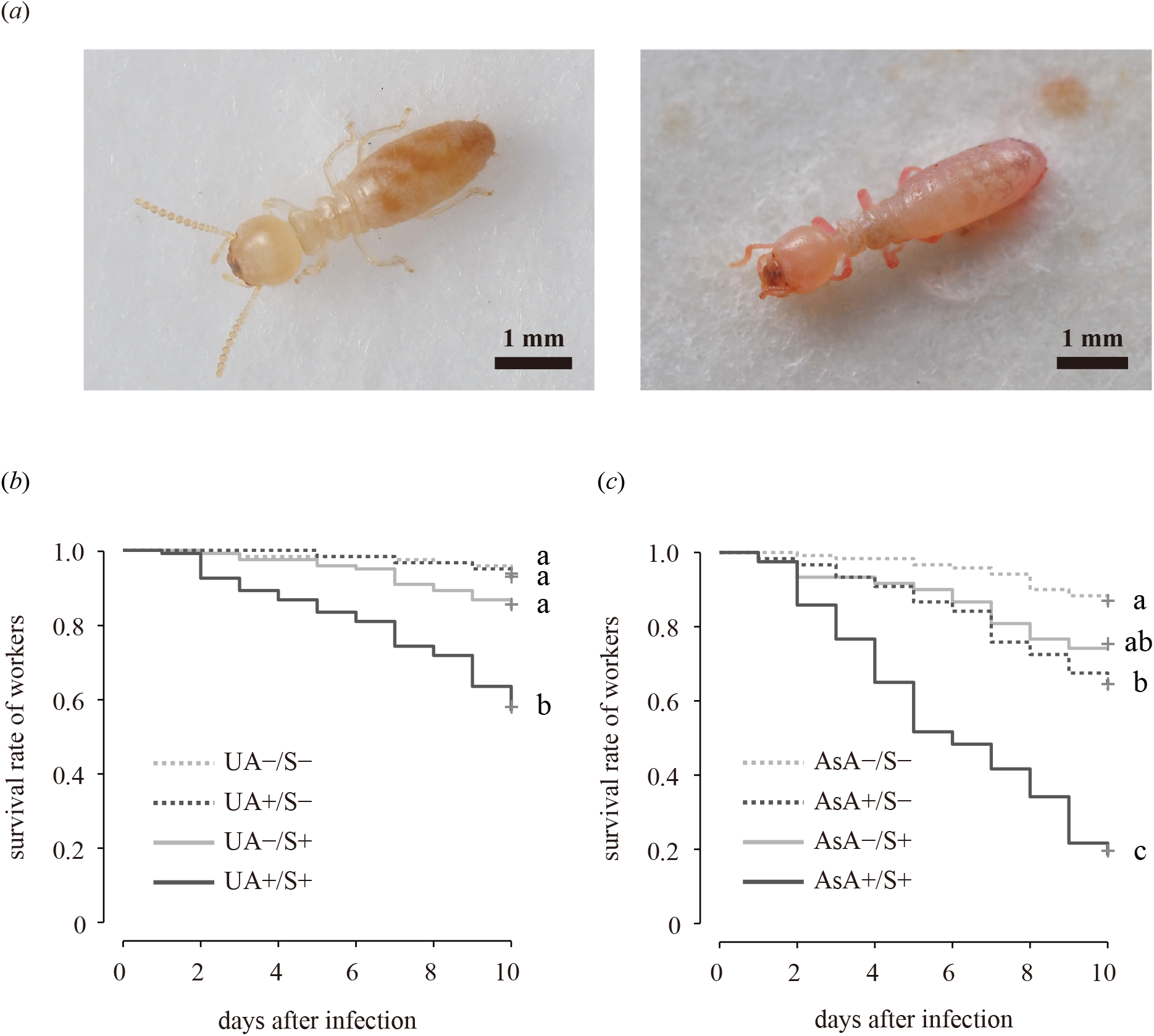
Infection assays using the opportunistic bacterium *Serratia marcescens*, which causes disease in immunocompromised termites. (*a*) Representative photo of the untreated worker (left panel) and dead termite bodies that have turned red due to opportunistic infection by *S. marcescens* (right panel). (*b*) Effects of uric acid administration on the survival rates of workers fed uric acid mixed in brown-rotted pinewood mixed cellulose (BPC) medium and those fed BPC medium only (‘UA+’ and ‘UA− ‘, respectively). (*c*) Effects of the administration of another antioxidant, ascorbic acid, on the survival rates of workers fed ascorbic acid mixed in BPC medium and those fed BPC medium only (‘AsA+’ and ‘AsA −’, respectively). ‘S+’ represents workers exposed to *S. marcescens* cultured in lysogeny broth (LB) liquid media and ‘S−’ means those exposed to LB liquid media only. Different letters indicate significant differences (log-rank test with Bonferroni correction, *p <* 0.008). (Online version in colour.)

## 4. Discussion

We showed that uric acid accumulation in workers after king replacement makes colonies susceptible to infectious diseases in the subterranean termite *R. speratus*. Our results can partially explain the proximate mechanisms involved in colony demise in this species. It is crucial for social insect colonies to reduce the risk of infection by pathogens, including opportunistic ones, because the high density, frequent contact and low genetic diversity in colonies are all expected to heighten the risk of pathogen transmission [44]. To compensate for this disadvantages, social insects exhibit not only individual immunity but also collective anti-pathogen activities called ‘social immunity’, such as allogrooming and carcass-burying [45–47]. Here, we point out that regulating the individual uric acid levels through the interactions among nestmates (e.g. transfer of nitrogen compounds) plays a pivotal role as a part of the social immunity in termites. While investigating the prevalence of this phenomenon among termite species (or social insect species, including honeybees) remains for future studies, it is known that workers accumulate uric acid inside fat bodies after maintenance under laboratory conditions (i.e. separated from kings) not only in *R. speratus* [23,48] but also in some other termite species [49–51]. In *R. speratus*, amounts of uric acid in worker bodies increase within three months after the removal of the primary king [55–57]. This rapid change suggests that uric acid accumulation in workers is independent of inbreeding depression after the emergence of secondary kings.

How does king replacement cause the accumulation of uric acid in worker bodies? Uric acid in termites has often been discussed in terms of nitrogen recycling [52–55]. Wood-eating termites feed on a nitrogen-poor diet [56–58] and have evolved efficient nitrogen acquisition and conservation strategies such as obtaining dietary nitrogen, nitrogen fixation by symbiotic microbes, and recycling of termite-derived products (e.g. nestmate bodies and exuviae) [59–64]. Additionally, termites do not void metabolites of acquired nitrogen compounds; instead, they are stored as uric acid, which is synthesised in fat bodies [49,65,66]. Uric acid is stored in specialised cells called urocytes of fat bodies in termites [46,65]. Intriguingly, Konishi et al. [47,66] showed that (i) only kings and queens have the enzyme to degrade uric acid, (ii) uric acid degradation contributes to reproduction and (iii) uric acid is provided by workers to the reproductive castes in *R. speratus*. Therefore, king replacement (loss of the primary kings) can alter the material flow within colonies, leading to uric acid accumulation in workers. Note that the increased amount of uric acid in worker bodies following the loss of the primary king does not return to normal after the appearance of the secondary king, which also can degrade uric acid [25–27]. Future studies should conduct a comparative analysis between primary kings and secondary kings in terms of aspects such as consumption of uric acid based on enzyme activity and influences on trophallaxis among colony members. It is also necessary to take into account the possibility that such uric acid accumulation in workers can be understood as not only stagnation of the material flow but also a response against oxidative stress caused by isolation from primary kings. Termites are already known to accumulate uric acid during stressful conditions [48,67]. In any case, although avoiding uric acid accumulation by workers after king replacement seems to be more adaptive, colonies depend on the presence of primary kings only, not secondary kings (and numerous queens). Further research is needed to determine whether termites cannot overcome the constraints or need not (colony death with loss of primary kings may have adaptive significance).

This study is the first to demonstrate the negative effect of uric acid on termites. The survival rate of workers with increased levels of uric acid was lower than those with normal uric acid levels when both were exposed to an opportunistic pathogen that causes disease to immunocompromised termites. Uric acid supplementation had no effect on the survival in the uninfected control, which indicates that uric acid itself does not have physical toxicity to termite tissues. Uric acid exerts a strong antioxidant activity against reactive oxygen species (ROS) [67], which damage biomolecules at high concentrations. However, appropriate amounts of ROS play a key role in the immune function of organisms [68]. ROS itself attacks pathogens in a similar manner as antimicrobial peptide (AMP) and melanin; moreover, they also act as a signalling molecule in the innate immune response. Our results show that the antioxidant activity of uric acid increases infectious disease risk through the depletion of ROS. Workers with high levels of uric acid had low ROS contents and survival rates against *Serratia* exposure. Similarly, low survival rates against *Serratia* infection were observed in workers fed another antioxidant. A similar phenomenon was reported in another insect species, the sand fly *Lutzomyia longipalpis*, where feeding uric acid as an exogenous ROS scavenger reduces survival after infection with *S. marcescens* [68]. Future studies should elucidate more detailed mechanisms that uric acid accumulation increases infectious disease risk in termites, such as by examining amounts of ROS and pathogens (*S. marcescens* and other bacteria and fungi) during infection. Furthermore, our findings also can lead to future technical innovations in applied entomology. Termites are one of the most economically important pests, which cause damage to crops and man-made structures estimated at over $50 billion per year in the world [69]. Our results offer a novel strategy for termite control, i.e. feeding termites excessive amounts of antioxidants to compromise termite immunity, thus accelerating infection and death from pathogens, including opportunistic pathogens. Antioxidants may provide an alternative to the toxic and expensive pesticides currently used worldwide.

In conclusion, the excessive accumulation of uric acid, a major product of nitrogen metabolism with antioxidant properties, increases the infectious disease risk in termites. Our results indicate that regulating the individual oxidant/antioxidant balance through the interactions among colony members is the key to the immunity of social insects. We provide further empirical evidence that social insect colonies act as a ‘superorganism’ that regulates metabolism at the social level [70,71]. Noteworthy, it is known that a high-protein diet reduces individual lifespan, leading to colony death in ants and honeybees [72,73]. Considering the physiological similarities in metabolism and immunity among insect species, we could apply our scenario to other social insects. This study opens new avenues for understanding the mechanisms underlying the maintenance of social systems in animals.

## Data accessibility

The dataset and R code supporting this article have been uploaded as part of the electronic supplementary materials.

## Author’s contributions

TK: conceptualisation, data curation, formal analysis, funding acquisition, project administration, investigation, methodology, resources, validation, visualisation, writing-original draft, writing-review and editing; MN: conceptualisation, data curation, formal analysis, investigation, methodology, resources, validation, visualisation, writing-review and editing; TN: conceptualisation, methodology, resources, visualisation, writing-review and editing; ET: conceptualisation, methodology, resources, visualisation, writing-review and editing; MT: conceptualisation, methodology, resources, visualisation, writing-review and editing; KM: conceptualisation, funding acquisition, project administration, resources, supervision, writing-review and editing. All authors gave final approval for publication and agreed to be held accountable for the work performed therein.

## Acknowledgements

We thank Tomoki Ishibashi, Shuya Nagai, Takehiro Morimoto, Chihiro Tamaki, Hiroki Noda, Hikaru Mitomo, Kiyotaka Yabe, and Tomohiro Nakazono for their assistance in collecting termites; Michihiko Takahashi and Matthew Kamiyama for helpful discussion. We also thank all other members of the Laboratory of Insect Ecology, Kyoto University for inspiring scientific discussions.

## References

1. Korb J. 2008 Termites, hemimetabolous diploid white ants? Front. Zool. 5, 1–9. (doi:10.1186/1742-9994-5-15)

2. Schultz TR. 2000 In search of ant ancestors. Proc. Natl. Acad. Sci. U. S. A. 97, 14028–14029. (doi:10.1073/pnas.011513798)

3. Bar-On YM, Phillips R, Milo R. 2018 The biomass distribution on Earth. Proc. Natl. Acad. Sci. U. S. A. 115, 6506–6511. (doi:10.1073/pnas.1711842115)

4. Wilson EO. 1971 The insect societies. Cambridge, MA: Harvard University Press.

5. Wilson EO. 1975 Sociobiology. Cambridge, MA: Harvard University Press.

6. Keller L, Genoud M. 1997 Extraordinary lifespans in ants: a test of evolutionary theories of ageing. Nature 389, 958–960. (doi:10.1038/40130)

7. Tasaki E, Takata M, Matsuura K. 2021 Why and how do termite kings and queens live so long? Philos. Trans. R. Soc. Lond. B Biol. Sci. 376, 20190740. (doi:10.1098/rstb.2019.0740)

8. Wyss-Huber M, Lüscher M. 1975 Protein synthesis in ‘fat body’ and ovary of the physogastric queen of Macrotermes subhyalinus. J. Insect Physiol. 21, 1697–1704. (doi:10.1016/0022-1910(75)90182-1)

9. Haydak MH. 1970 Honey bee nutrition. Annu. Rev. Entomol. 15, 143–156. (doi:10.1146/annurev.en.15.010170.001043)

10. Myles TG. 1999 Review of secondary reproduction in termites (Insecta: Isoptera) with comments on its role in termite ecology and social evolution. Sociobiology 33, 1–43.

11. Oster GF, Wilson EO. 1978 Caste and ecology in the social insects. Princeton, NJ: Princeton University Press.

12. Lepage M, Darlington JPEC. 2000 Population dynamics of termites. In Termites: evolution, sociality, symbioses, ecology (eds T Abe, DE Bignell, M Higashi), pp. 333–361. Dordrecht: Springer Netherlands. (doi:10.1007/978-94-017-3223-9_16)

13. Sun Q, Haynes KF, Zhou X. 2018 Managing the risks and rewards of death in eusocial insects. Philos. Trans. R. Soc. Lond. B Biol. Sci. 373, 20170258.

14. Collins NM. 1981 Populations, age structure and survivorship of colonies of Macrotermes bellicosus (Isoptera: Macrotermitinae). J. Anim. Ecol. 50, 293–312. (doi:10.2307/4046)

15. Chouvenc T, Ban PM, Su N-Y. 2022 Life and death of termite colonies, a decades-long age demography perspective. Front. Ecol. Evol. 10, 911042. (doi:10.3389/fevo.2022.911042)

16. Ellis JD, Evans JD, Pettis J. 2010 Colony losses, managed colony population decline, and Colony Collapse Disorder in the United States. J. Apic. Res. 49, 134–136. (doi:10.3896/ibra.1.49.1.30)

17. vanEngelsdorp D et al. 2009 Colony collapse disorder: a descriptive study. PLoS One 4, e6481. (doi:10.1371/journal.pone.0006481)

18. Dainat B, Evans JD, Chen YP, Gauthier L, Neumann P. 2012 Predictive markers of honey bee colony collapse. PLoS One 7, e32151. (doi:10.1371/journal.pone.0032151)

19. Chouvenc T, Bardunias P, Efstathion CA, Chakrabarti S, Elliott ML, Giblin-Davis R, Su N-Y. 2013 Resource opportunities from the nest of dying subterranean termite (Isoptera: Rhinotermitidae) colonies: a laboratory case of ecological succession. Ann. Entomol. Soc. Am. 106, 771–778. (doi:10.1603/an13104)

20. Matsuura K. 2017 Evolution of the asexual queen succession system and its underlying mechanisms in termites. J. Exp. Biol. 220, 63–72. (doi:10.1242/jeb.142547)

21. Matsuura K, Vargo EL, Kawatsu K, Labadie PE, Nakano H, Yashiro T, Tsuji K. 2009 Queen succession through asexual reproduction in termites. Science 323, 1687. (doi:10.1126/science.1169702)

22. Matsuura K, Mizumoto N, Kobayashi K, Nozaki T, Fujita T, Yashiro T, Fuchikawa T, Mitaka Y, Vargo EL. 2018 A genomic imprinting model of termite caste determination: not genetic but epigenetic inheritance influences offspring caste fate. Am. Nat. 191, 677–690. (doi:10.1086/697238)

23. Konishi T, Tasaki E, Takata M, Matsuura K. 2023 King- and queen-specific degradation of uric acid contributes to reproduction in termites. Proc. Biol. Sci. 290, 20221942. (doi:10.1098/rspb.2022.1942)

24. Becker BF, Reinholz N, Leipert B, Raschke P, Permanetter B, Gerlach E. 1991 Role of uric acid as an endogenous radical scavenger and antioxidant. Chest 100, 176S–181S. (doi:10.1378/chest.100.3_supplement.176s)

25. Dowling DK, Simmons LW. 2009 Reactive oxygen species as universal constraints in life-history evolution. Proc. Biol. Sci. 276, 1737–1745. (doi:10.1098/rspb.2008.1791)

26. Apel K, Hirt H. 2004 Reactive oxygen species: metabolism, oxidative stress, and signal transduction. Annu. Rev. Plant Biol. 55, 373–399. (doi:10.1146/annurev.arplant.55.031903.141701)

27. Dröge W. 2002 Free radicals in the physiological control of cell function. Physiol. Rev. 82, 47–95. (doi:10.1152/physrev.00018.2001)

28. Hayashi Y, Kitade O, Kojima J-I. 2003 Parthenogenetic reproduction in neotenics of the subterranean termite Reticulitermes speratus (Isoptera: Rhinotermitidae). Entomol. Sci. 6, 253–257. (doi:10.1046/j.1343-8786.2003.00030.x)

29. Mitaka Y, Akino T, Matsuura K. 2023 Development of a standard medium for culturing the termite Reticulitermes speratus. Insectes Soc. 70, 265–274. (doi:10.1007/s00040-023-00907-6)

30. Elliott KL, Stay B. 2007 Juvenile hormone synthesis as related to egg development in neotenic reproductives of the termite Reticulitermes flavipes, with observations on urates in the fat body. Gen. Comp. Endocrinol. 152, 102–110. (doi:10.1016/j.ygcen.2007.03.003)

31. Costa-Leonardo AM, Laranjo LT, Janei V, Haifig I. 2013 The fat body of termites: functions and stored materials. J. Insect Physiol. 59, 577–587. (doi:10.1016/j.jinsphys.2013.03.009)

32. Nozaki T, Tasaki E, Matsuura K. 2023 Cell type specific polyploidization in the royal fat body of termite queens. Zoological Lett 9, 20. (doi:10.1186/s40851-023-00217-6)

33. Grimont PA, Grimont F. 1978 The genus Serratia. Annu. Rev. Microbiol. 32, 221–248. (doi:10.1146/annurev.mi.32.100178.001253)

34. Osbrink WLA, Williams KS, Connick WJ, Wright MS, Lax AR. 2001 Virulence of bacteria associated with the Formosan subterranean termite (Isoptera: Rhinotermitidae) in New Orleans, LA. Environ. Entomol. 30, 443–448. (doi:10.1603/0046-225X-30.2.443)

35. Connick WJ, Osbrink WLA, Wright MS, Williams KS, Daigle DJ, Boykin DL, Lax AR. 2001 Increased mortality of Coptotermes formosanus (Isoptera: Rhinotermitidae) exposed to eicosanoid biosynthesis inhibitors and Serratia marcescens (Eubacteriales: Enterobacteriaceae). Environ. Entomol. 30, 449–455. (doi:10.1603/0046-225X-30.2.449)

36. De Bach PH, Mcomie WA. 1939 New diseases of termites caused by bacteria. Ann. Entomol. Soc. Am. 32, 137–146. (doi:10.1093/aesa/32.1.137)

37. Inagaki T, Matsuura K. 2018 Extended mutualism between termites and gut microbes: nutritional symbionts contribute to nest hygiene. Naturwissenschaften 105, 52. (doi:10.1007/s00114-018-1580-y)

38. Lehejčkova R, Demnerova K, Kralova B, Лехейчкова Р, демнерова К, ралова В, Лехейчкова Р, демнерова К, Кралова В. 1987 Screening of microorganisms with uricase activity. Biotechnology & Bioindustry 2, 25–27. (doi:10.1080/02052067.1987.10819296)

39. Pichon A, Kutnik M, Leniaud L, Darrouzet E, Châline N, Dupont S, Bagnères A-G. 2007 Development of experimentally orphaned termite worker colonies of two Reticulitermes species (Isoptera: Rhinotermitidae). Sociobiology 50, 1015–1034.

40. Ghesini S, Marini M. 2009 Caste differentiation and growth of laboratory colonies of Reticulitermes urbis (Isoptera, Rhinotermitidae). Insectes Soc. 56, 309–318. (doi:10.1007/s00040-009-0025-1)

41. Wilson K. 2001 Preparation of genomic DNA from bacteria. Curr. Protoc. Mol. Biol. Chapter 2, Unit 2.4. (doi:10.1002/0471142727.mb0204s56)

42. Altschul SF, Madden TL, Schäffer AA, Zhang J, Zhang Z, Miller W, Lipman DJ. 1997 Gapped BLAST and PSI-BLAST: a new generation of protein database search programs. Nucleic Acids Res. 25, 3389–3402. (doi:10.1093/nar/25.17.3389)

43. R Core Team. 2024 R: a language and environment for statistical computing. Vienna, Austria: R Foundation for Statistical Computing.

44. Schmid-Hempel P. 1998 Parasites in social insects. Princeton, NJ: Princeton University Press.

45. Cremer S, Armitage SAO, Schmid-Hempel P. 2007 Social immunity. Curr. Biol. 17, R693–R702. (doi:10.1016/j.cub.2007.06.008)

46. Wilson-Rich N, Spivak M, Fefferman NH, Starks PT. 2009 Genetic, individual, and group facilitation of disease resistance in insect societies. Annu. Rev. Entomol. 54, 405–423. (doi:10.1146/annurev.ento.53.103106.093301)

47. Cremer S, Sixt M. 2009 Analogies in the evolution of individual and social immunity. Philos. Trans. R. Soc. Lond. B Biol. Sci. 364, 129–142. (doi:10.1098/rstb.2008.0166)

48. Tasaki E, Sakurai H, Nitao M, Matsuura K, Iuchi Y. 2017 Uric acid, an important antioxidant contributing to survival in termites. PLoS One 12, e0179426. (doi:10.1371/journal.pone.0179426)

49. Potrikus CJ, Breznak JA. 1980 Uric acid in wood-eating termites. Insect Biochem. 10, 19–27. (doi:10.1016/0020-1790(80)90034-7)

50. Lovelock M, O’Brien RW, Slaytor M. 1985 Effect of laboratory containment on the nitrogen metabolism of termites. Insect Biochem. 15, 503–509. (doi:10.1016/0020-1790(85)90063-0)

51. Chappell DJ, Slaytor M. 1993 Uric acid synthesis in freshly collected and laboratory-maintained Nasutitermes walkeri hill. Insect Biochem. Mol. Biol. 23, 499–506. (doi:10.1016/0965-1748(93)90058-Z)

52. Potrikus CJ, Breznak JA. 1980 Anaerobic degradation of uric Acid by gut bacteria of termites. Appl. Environ. Microbiol. 40, 125–132. (doi:10.1128/aem.40.1.125-132.1980)

53. Potrikus CJ, Breznak JA. 1981 Gut bacteria recycle uric acid nitrogen in termites: A strategy for nutrient conservation. Proc. Natl. Acad. Sci. U. S. A. 78, 4601–4605. (doi:10.1073/pnas.78.7.4601)

54. Thong-On A, Suzuki K, Noda S, Inoue J-I, Kajiwara S, Ohkuma M. 2012 Isolation and characterization of anaerobic bacteria for symbiotic recycling of uric acid nitrogen in the gut of various termites. Microbes Environ. 27, 186–192. (doi:10.1264/jsme2.me11325)

55. Waidele L, Korb J, Voolstra CR, Dedeine F, Staubach F. 2019 Ecological specificity of the metagenome in a set of lower termite species supports contribution of the microbiome to adaptation of the host. Anim Microbiome 1, 13. (doi:10.1186/s42523-019-0014-2)

56. La Fage JP, Nutting WL. 1978 Nutrient dynamics of termites. In Production ecology of ants and termites (ed MV Brian), pp. 165–232. Cambridge, UK: Cambridge University Press.

57. Higashi M, Abe T, Burns TP. 1992 Carbon-nitrogen balance and termite ecology. Proc. Biol. Sci. 249, 303–308. (doi:10.1098/rspb.1992.0119)

58. Matsumoto T. 1976 The role of termites in an equatorial rain forest ecosystem of West Malaysia. Oecologia 22, 153–178. (doi:10.1007/BF00344714)

59. Hungate RE. 1941 Experiments on the nitrogen economy of termites. Ann. Entomol. Soc. Am. 34, 467–489. (doi:10.1093/aesa/34.2.467)

60. Benemann JR. 1973 Nitrogen fixation in termites. Science 181, 164–165. (doi:10.1126/science.181.4095.164)

61. Breznak JA, Brill WJ, Mertins JW, Coppel HC. 1973 Nitrogen fixation in termites. Nature 244, 577–580. (doi:10.1038/244577a0)

62. Chouvenc T. 2020 Limited survival strategy in starving subterranean termite colonies. Insectes Soc. 67, 71–82. (doi:10.1007/s00040-019-00729-5)

63. Tong RL, Aguilera-Olivares D, Chouvenc T, Su N-Y. 2021 Nitrogen content of the exuviae of Coptotermes gestroi (Wasmann) (Blattodea: Rhinotermitidae). Heliyon 7, e06697. (doi:10.1016/j.heliyon.2021.e06697)

64. Mullins A, Chouvenc T, Su N-Y. 2021 Soil organic matter is essential for colony growth in subterranean termites. Sci. Rep. 11, 21252. (doi:10.1038/s41598-021-00674-z)

65. Breznak JA. 1982 Intestinal microbiota of termites and other xylophagous insects. Annu. Rev. Microbiol. 36, 323–323. (doi:10.1146/annurev.mi.36.100182.001543)

66. Brune A. 2014 Symbiotic digestion of lignocellulose in termite guts. Nat. Rev. Microbiol. 12, 168–180. (doi:10.1038/nrmicro3182)

67. Elsner D, Hartfelder K, Korb J. 2021 Molecular underpinnings of division of labour among workers in a socially complex termite. Sci. Rep. 11, 18269. (doi:10.1038/s41598-021-97515-w)

68. Diaz-Albiter H, Sant’Anna MRV, Genta FA, Dillon RJ. 2012 Reactive oxygen species-mediated immunity against Leishmania mexicana and Serratia marcescens in the phlebotomine sand fly Lutzomyia longipalpis. J. Biol. Chem. 287, 23995–24003. (doi:10.1074/jbc.M112.376095)

69. Korb J. 2007 Termites. Curr. Biol. 17, R995–9. (doi:10.1016/j.cub.2007.10.033)

70. Friedman DA, Johnson BR, Linksvayer TA. 2020 Distributed physiology and the molecular basis of social life in eusocial insects. Horm. Behav. 122, 104757. (doi:10.1016/j.yhbeh.2020.104757)

71. Negroni MA, LeBoeuf AC. 2023 Metabolic division of labor in social insects. Curr Opin Insect Sci 59, 101085. (doi:10.1016/j.cois.2023.101085)

72. Pirk CWW, Boodhoo C, Human H, Nicolson SW. 2010 The importance of protein type and protein to carbohydrate ratio for survival and ovarian activation of caged honeybees (Apis mellifera scutellata). Apidologie 41, 62–72. (doi:10.1051/apido/2009055)

73. Dussutour A, Simpson SJ. 2012 Ant workers die young and colonies collapse when fed a high-protein diet. Proc. Biol. Sci. 279, 2402–2408. (doi:10.1098/rspb.2012.0051)

