## Supplementary materials for "What makes a society dead: accumulation of uric acid increases infectious disease risk in termites"

**This file includes:**

Figure S1

**Other Supplementary Materials for this manuscript include the following:**

Dataset S1

R code S1

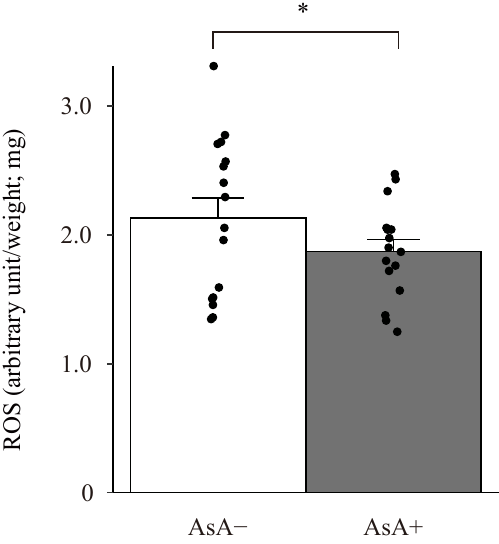

**Figure S1.** Effects of the administration of another antioxidant, ascorbic acid, on reactive oxygen species (ROS) in workers fed ascorbic acid mixed in brown-rotted pinewood mixed cellulose (BPC) medium and those fed BPC medium only (‘AsA+’ and ‘AsA−’, respectively). Black points show individual termites, jittered to reduce overlap between points. Error bars denote the standard error of the mean (SEM). Asterisks indicate significant differences (likelihood ratio test, **p* < 0.05, ***p* < 0.01, ****p* < 0.001).
